# Inflammatory cytokines mediate the induction of and awakening from metastatic dormancy

**DOI:** 10.1101/2024.08.05.606588

**Authors:** Paulo Pereira, Joshua Panier, Marc Nater, Michael Herbst, Anna Laura Calvanese, Hakan Köksal, Héctor Castañón, Virginia Cecconi, Paulino Tallón de Lara, Steve Pascolo, Maries van den Broek

**Affiliations:** Institute of Experimental Immunology, University of Zurich, Switzerland; Department of Dermatology, University Hospital Zurich, Switzerland; Division of Cancer Medicine, The University of Texas MD Anderson Cancer Center, Houston, Texas, USA

**Keywords:** Metastatic dormancy, IFNγ, IL-17A, breast cancer

## Abstract

Metastases arise from disseminated cancer cells (DCCs) that detach from the primary tumor and seed distant organs. There, quiescent DCCs can survive for an extended time, a state referred to as metastatic dormancy. The mechanisms governing the induction, maintenance, and awakening from metastatic dormancy are unclear. We show that the differentiation of dormancy-inducing CD8^+^ T cells requires CD4^+^ T cell help, and that IFNγ directly induces dormancy in DCCs. The maintenance of metastatic dormancy, however, is independent of T cells. Instead, awakening from dormancy requires an inflammatory signal, and we identified CD4^+^ T cell-derived IL- 17A as an essential wake-up signal for dormant DCCs in the lungs.

Thus, the induction and awakening from metastatic dormancy require an external stimulus, while the maintenance of dormancy does not rely on the continuous surveillance by lymphocytes.

## INTRODUCTION

Most of the research in the field of metastasis focuses on aggressive and rapidly metastasizing cancers. In some human cancers, however, the period between resection of the primary tumor and the appearance of macroscopic metastases can be as long as decades.^1–3^ The relevance of this fact is illustrated by the knowledge that approximately two-thirds of breast cancer-related deaths occur more than 5 years after the removal of the primary tumor.^4^

Those late metastatic relapses are thought to originate from cancer cells that remain quiescent for a long time. This is supported by the finding that disseminated cancer cells (DCCs) are present in a dormant state in the bone marrow of early-stage breast cancer patients.^5,6^ Moreover, several studies showed that patients who received organs from individuals who had been cured of non-invasive melanoma decades ago can develop melanomas from donor origin,^7^ highlighting the longevity of these quiescent cancer cells.

Although the mechanisms governing dormancy and awakening are largely unknown, several factors that influence the duration of dormancy have been suggested. The tyrosine kinase Src is required for the survival of dormant breast cancer cells,^8^ and the transcription factor ZFP281 keeps early DCCs in the lungs in a dormant state.^9^ In human carcinoma cell lines, a high p38/ERK ratio is associated with cell-cycle arrest, while a low p38/ERK ratio is observed in proliferating cells.^10^ *In vivo* studies showed that the orphan nuclear receptor NR2F1, also required for the induction of dormancy, is upregulated in dormant head and neck squamous carcinoma and in disseminated prostate cancer cells in individuals with dormant disease.^11^

Despite the importance of cancer cell-intrinsic mechanisms, extracellular signals from the microenvironment also regulate metastatic dormancy. Wnt5a, a protein paradoxically described to have oncogenic and tumor suppressive functions,^12^ induces metastatic dormancy in prostate cancer cells in the lungs,^13^ and the bone morphogenic protein 7 induces cell-cycle arrest in prostate cancer stem-like cells in the bone^14^. Similarly, osteopontin stops the proliferation of acute lymphoblastic leukemia cells in the bone marrow, protecting them from chemotherapeutic interventions.^15^

Nevertheless, despite their clinical relevance, the pathways responsible for activating these mechanisms remain unidentified. Even less clear are the pathways and cells responsible for the awakening from metastatic dormancy.

Some factors, such as tobacco-induced neutrophil extracellular traps, can awaken dormant cancer cells in the lungs,^16^ but the role of other immune cells and cytokines in the awakening of dormant cancer cells remains unknown.

Here, we identify the role of the adaptive immunity in the production of the cytokines required for the induction and awakening from metastatic dormancy, whereas the maintenance of metastatic dormancy is independent of T cell surveillance.

## RESULTS

### Induction of metastatic dormancy requires CD4^+^ T cell help

We used a preclinical breast cancer model, 4T07, for metastatic dormancy. In this model, cancer cells disseminate from the primary tumor to the lungs, where they persist as quiescent cells.^17^ We recently showed that the induction of metastatic dormancy in the lungs is mediated by CD8^+^ PD-1^+^ CD39^+^ T cells that produce TNFα and IFNγ.^18,19^ The generation and maintenance of protective CD8^+^ T cells in the context of cancer relies on CD4^+^ T cells.^20^ To study the contribution of CD4^+^ T cells in the induction of metastatic dormancy, we depleted CD4^+^ T cells before and during the priming of CD8^+^ T cells by the antigens derived from the primary 4T07-LZ tumor (expressing luciferase and ZsGreen). We then quantified the lung metastatic load 25 days later (Figure 1A). Mice lacking CD4^+^ T cells had a significantly higher metastatic burden in the lungs than non-depleted mice (Figures 1B and 1C), suggesting that CD4^+^ T cells are required for the induction of metastatic dormancy. To investigate whether CD4^+^ T cells induce dormancy by promoting the development of the dormancy-inducing CD8^+^ T cells, we characterized the intratumoral CD8^+^ T cells in CD4-depleted (unhelped CD8^+^ T cells) and non-depleted (helped CD8^+^ T cells) mice by high-dimensional flow cytometry (Figures 1D and 1E). Whereas the frequency of the identified 8 clusters was similar in both experimental conditions, unhelped CD8^+^ T cells were nearly bereft of IFNγ^+^ CD8^+^ T effector cells (cluster 1) (Figure 1F). To determine whether intratumoral CD4^+^ T cells, besides providing help to CD8^+^ T cells, directly protect against metastases, we intravenously injected intratumoral CD4^+^, helped CD8^+^ or unhelped CD8^+^ T cells together with 4T07-LZ cells into naïve mice, and measured their protective capacity by IVIS (Figure S1A). Donor mice lacking CD4^+^ T cells had a significantly larger tumor size (Figure S1B) and tumor weight (Figure S1C) than isotype-treated donor mice. We quantified the metastatic burden in the lungs of the recipient mice at days 7 and 18, assessing the protection conferred by T cells against the progression of metastases (Figure S1D). Three out of four mice adoptively transferred with helped CD8^+^ T cells displayed a lower median fold change than control mice, opposed to 1 out of 4 mice adoptively transferred with CD4^+^ T cells (Figure S1D). These findings suggest that CD4^+^ T cells can’t control the proliferation of cancer cells.

**Figure 1.**
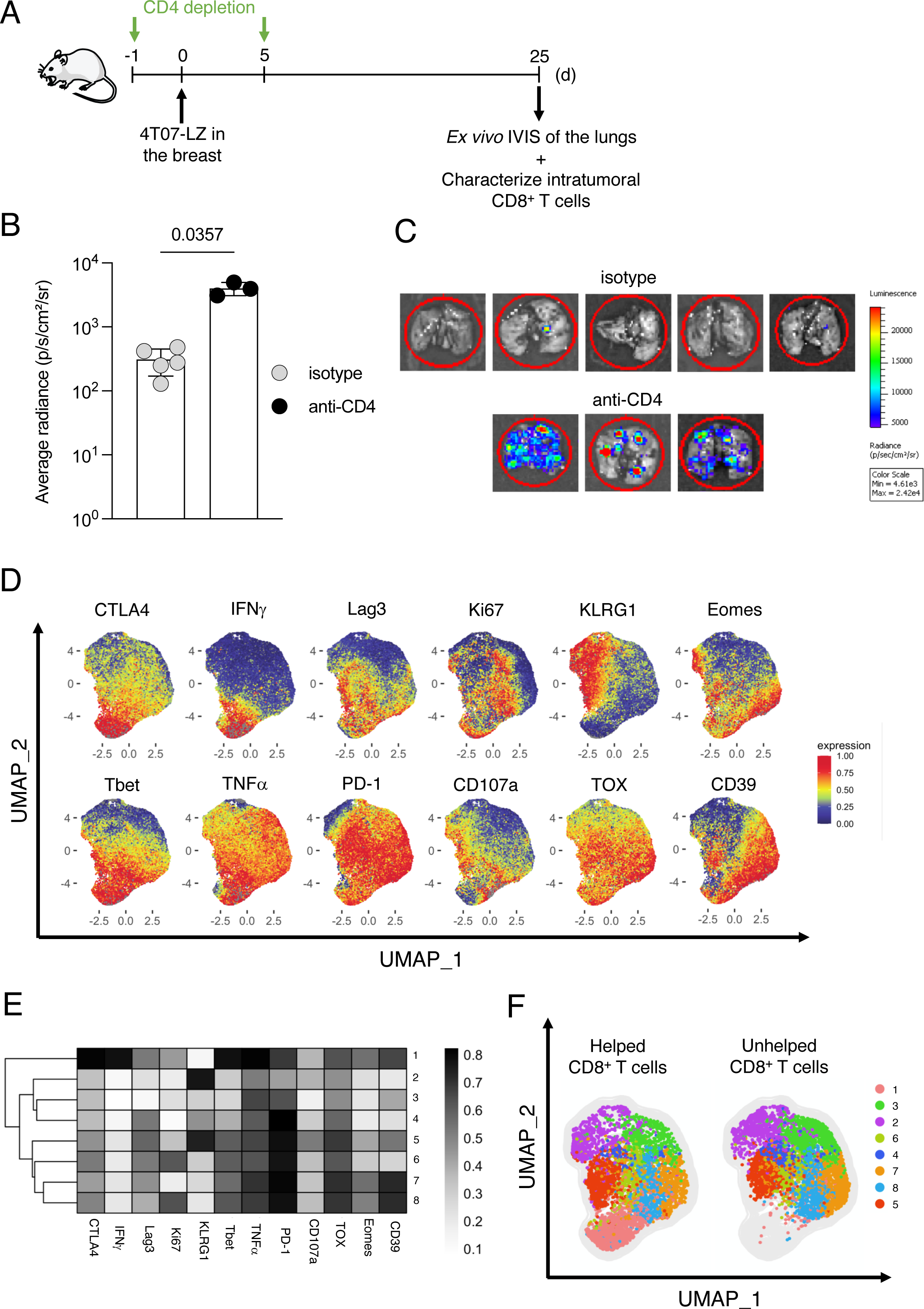
Induction of metastatic dormancy requires CD4^+^ T cell help. **(A)** Experimental design. Female BALB/cJRj mice received 0.5 mg isotype or anti- CD4 antibody i.p. at days -1 and +5 relative to the injection of 10^5^ 4T07-LZ cells in the mammary fat pad. The lungs and the primary tumor were collected on day 25 to quantify the metastatic burden and characterize intratumoral CD8^+^ T cells. **(B)** Quantification of the bioluminescent signal from the lungs. Isotype, n = 5; anti- CD4, n = 3. Statistical analysis was performed with a Mann-Whitney test. Each symbol represents an individual mouse. Data are shown as the mean ± standard deviation. The experiment is a representative example of 2 independent experiments. **(C)** Bioluminescence images of the lungs on day 25. **(D)** UMAP visualization of cytokines, proteins and markers after gating on single, live, CD45^+^, CD11b^-^, CD3^+^, CD8^+^ cells. Isotype, n = 5, anti-CD4, n = 3. **(E)** Heatmap with the relative expression of the tested markers in the identified clusters. **(F)** UMAP visualization of the clusters of intratumoral CD8^+^ T cells.

Altogether, these findings suggest that CD4^+^ T cells do not directly induce dormancy in cancer cells, but rather help in the development of the dormancy-inducing CD8^+^ T cells.

### Sensing of IFNγ by cancer cells induces metastatic dormancy

To determine whether IFNγ acts directly on cancer cells or indirectly via other cells, we generated IFNγR1-deficient 4T07-LZ cells (Figure S2A). To functionally confirm IFNγR1 deficiency, we analyzed the IFNγ-induced expression of PD-L1 on control 4T07-LZ*^nt^* (non-targeting-sgRNA) and 4T07-LZ^Δ*Ifngr*1^. Our observation that IFNγ only induced the expression of PD-L1 in the control cell line confirmed the successful deletion of *Ifngr1* (Figure S2B). We then orthotopically injected 4T07-LZ*^nt^* or 4T07- LZ^Δ*Ifngr*1^ cells, monitored tumor growth, and screened the lungs for metastatic outbreaks 21 days later (Figure 2A). In contrast to control tumors, most IFNγR1- deficient tumors started decreasing in size from day 14 onwards, and several tumors had regressed completely by day 21 (Figure 2B).

**Figure 2.**
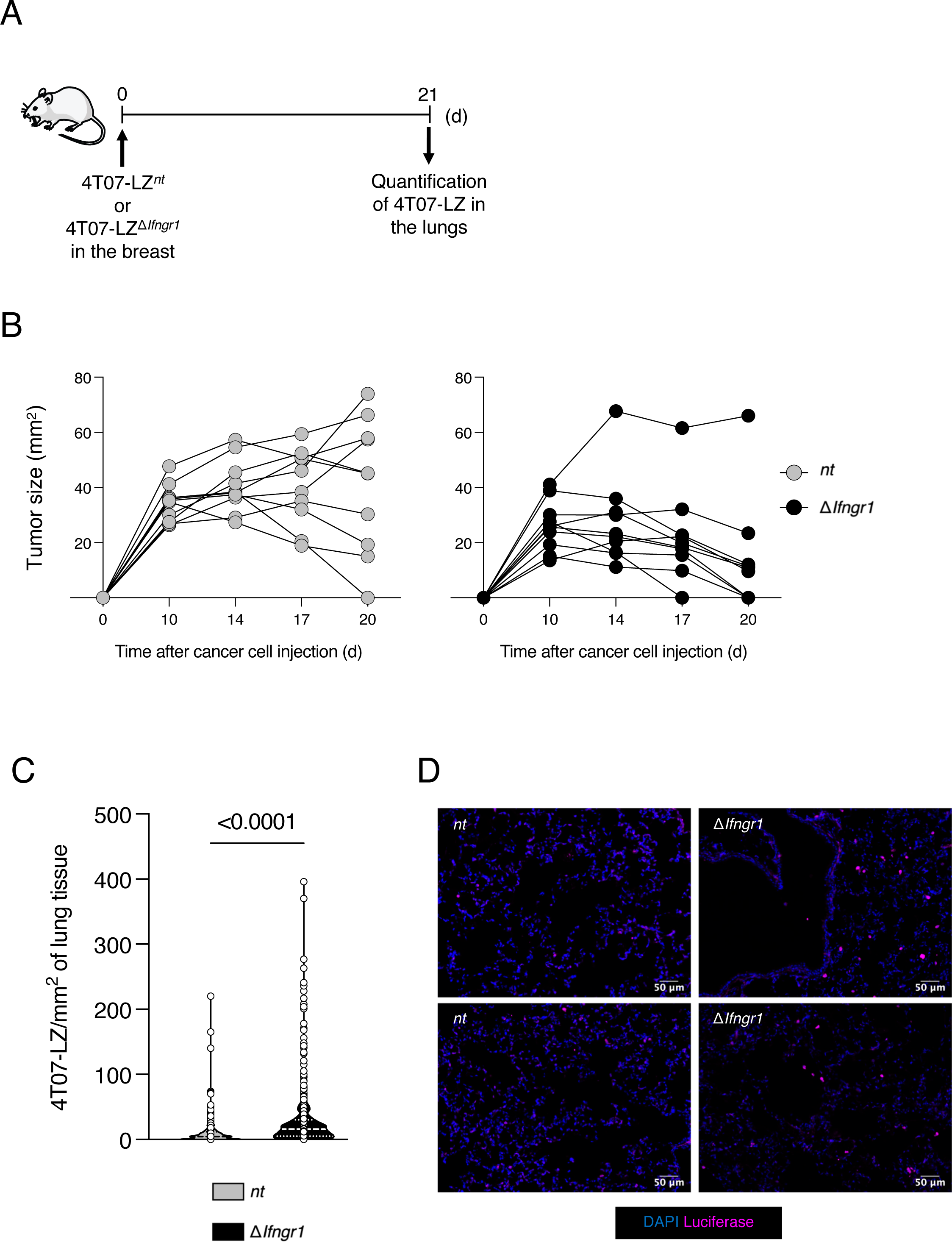
Sensing of IFNγ by cancer cells induces metastatic dormancy. **(A)** Experimental design. 4T07-mCh*^nt^* = 10; 4T07-mCh^Δ*Ifngr*1^ = 10. **(B)** Growth curves of the primary tumor from day 10 until the endpoint. Each symbol represents the tumor size of a single mouse. **(C)** Quantification of 4T07-LZ in the lungs on day 21. Statistical analysis was performed with a Mann-Whitney test. Each dot represents the number of cancer cells in a randomly chosen location in the lungs. The quartiles are represented by the dotted lines, the median is represented by the dashed line. **(D)** Immunofluorescence images of the lungs on day 21.

The failure to upregulate PD-L1 in response to IFNγ may explain the reduced growth of 4T07-LZ^Δ*Ifngr*1^ primary tumors *in vivo*. Indeed, 4T07-mCh^Δ*Ifngr*1^ cells expressed significantly less PD-L1 *in vivo* than 4T07-mCh*^nt^* cells (Figure S2C), which may result in better immune control and consequent decrease in tumor size.

Irrespective of the smaller tumor size, the lungs of mice bearing 4T07-LZ^Δ*Ifngr*1^ tumors contained significantly more DCCs than those of mice with 4T07-LZ*^nt^* tumors (Figures 2C and 2D), suggesting that sensing of IFNγ by DCCs contributes to dormancy. This is further supported by our observation that CD8^+^ T cells in IFNγR1-deficient and control 4T07-mCh tumors are phenotypically similar (Figures S2D and S2E).

Altogether, we show that IFNγ is the main driver of CD8^+^ T cell-induced metastatic dormancy of breast cancer.

### T and NK cells are dispensable for the maintenance of metastatic dormancy

After having identified the essential role of T cells in the induction of metastatic dormancy, we evaluated their role in maintaining dormancy. Therefore, we depleted T cells and analyzed the lungs for metastatic outbreaks. Before depletion, we resected the primary tumor to prevent the continuous influx of DCCs into the lungs (Figure 3A). To confirm that dormant DCCs persist in the lungs after the resection of the primary tumor without any further intervention, we screened the lungs for cancer cells in the lungs 63 days after resection (Figure S3A). We did not observe any nodules on the surface of the lungs or spontaneous awakening of the DCCs, but we observed single, non-proliferating cancer cells by immunofluorescence (Figures S3B and S3C).

**Figure 3.**
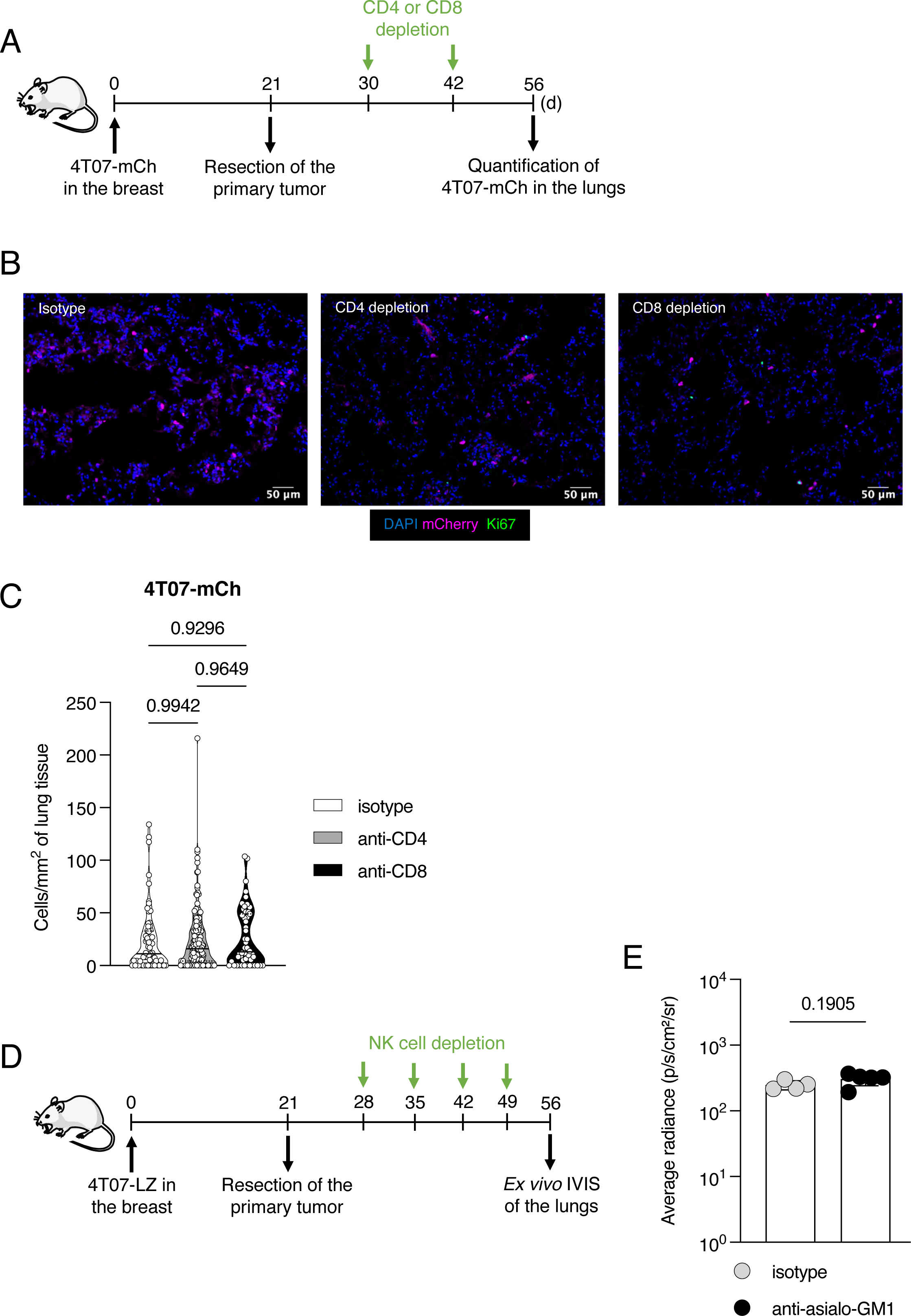
T and NK cells are dispensable for the maintenance of metastatic dormancy. **(A)** Experimental design. 4T07-mCh cells (10^5^) were injected into the mammary fat pad and the primary tumor was resected on day 21. Mice received 0.5 mg of isotype, anti-CD4, or anti-CD8 antibodies i.p. on days 30 and 42. The lungs were collected on day 56 to quantify the cancer cells by immunofluorescence. Isotype, n = 4; anti-CD4, n = 5; anti-CD8, n = 3. **(B)** Immunofluorescence images of the lungs on day 56. The experiment is a representative example of 2 independent experiments. **(C)** Quantification of the DCCs in the lungs on day 56. Each dot represents the number of cancer cells in a randomly chosen location in the lungs. Statistical analyses were performed with Brown-Forsythe and Welch ANOVA tests with a Dunnett’s T3 multiple comparisons test. **(D)** Experimental design. Mice received 50 µg of isotype or anti-asialo-GM1 antibodies i.p. once per week, starting one week after tumor resection until the endpoint. On day 56, the lungs were collected for quantification of the metastatic load by IVIS. Isotype, n = 4; anti-aisalo-GM1, n = 5. **(E)** Quantification of the bioluminescent signal from the lungs. Isotype = 4; anti-asialo- GM-1 = 5. Statistical analysis was performed with a Mann-Whitney test. Each symbol represents an individual mouse. Data are shown as the mean ± standard deviation.

The depletion of CD4^+^ or CD8^+^ T cells did not result in metastatic outbreaks or increase the number of DCCs in the lungs (Figures 3B and 3C), nor did the simultaneous depletion of CD4^+^ and CD8^+^ T cells (Figures S4A-C). The depletion of T cells was confirmed by flow cytometry on the blood (Figure S4D).

Because NK cells are also important producers of IFNγ, we depleted NK cells for 4 weeks after the resection of the primary tumor, after which we screened the lungs for metastatic outbreaks (Figure 3D). The depletion of NK cells did not increase the metastatic load in the lungs, compared to isotype-treated mice (Figure 3E), suggesting that NK cells are also not required for the maintenance of the dormant phenotype in the DCCs.

Altogether, our data show that NK and T cells are dispensable for the maintenance of metastatic dormancy.

### LPS-induced inflammation awakens dormant cancer cells

Based on our findings, we proposed that dormant DCCs may not require continuous signals to remain dormant but rather stimuli to awaken, such as inflammation. We used lipopolysaccharide (LPS), a well-characterized inflammatory stimulus, to test this hypothesis. Specifically, we repeatedly administered LPS intranasally (i.n.) after resecting the primary tumor and analyzed the lungs for metastatic outbreaks 12 hours and 7 days (168 hours) after the last LPS administration (E12 and E168, respectively) (Figure 4A). On E12, we observed a trend towards more DCCs in the lungs of LPS-treated mice, when compared to the PBS-treated controls (Figures 4B and 4C). On E168, the number of DCCs in the lungs of LPS-treated mice was significantly higher than in control mice (Figures 4B and 4C). The observation that a higher proportion of DCCs is cycling in the lungs of LPS-treated mice on E12 (Figure 4C) suggests a proliferative burst that is responsible for the observed awakening.

**Figure 4.**
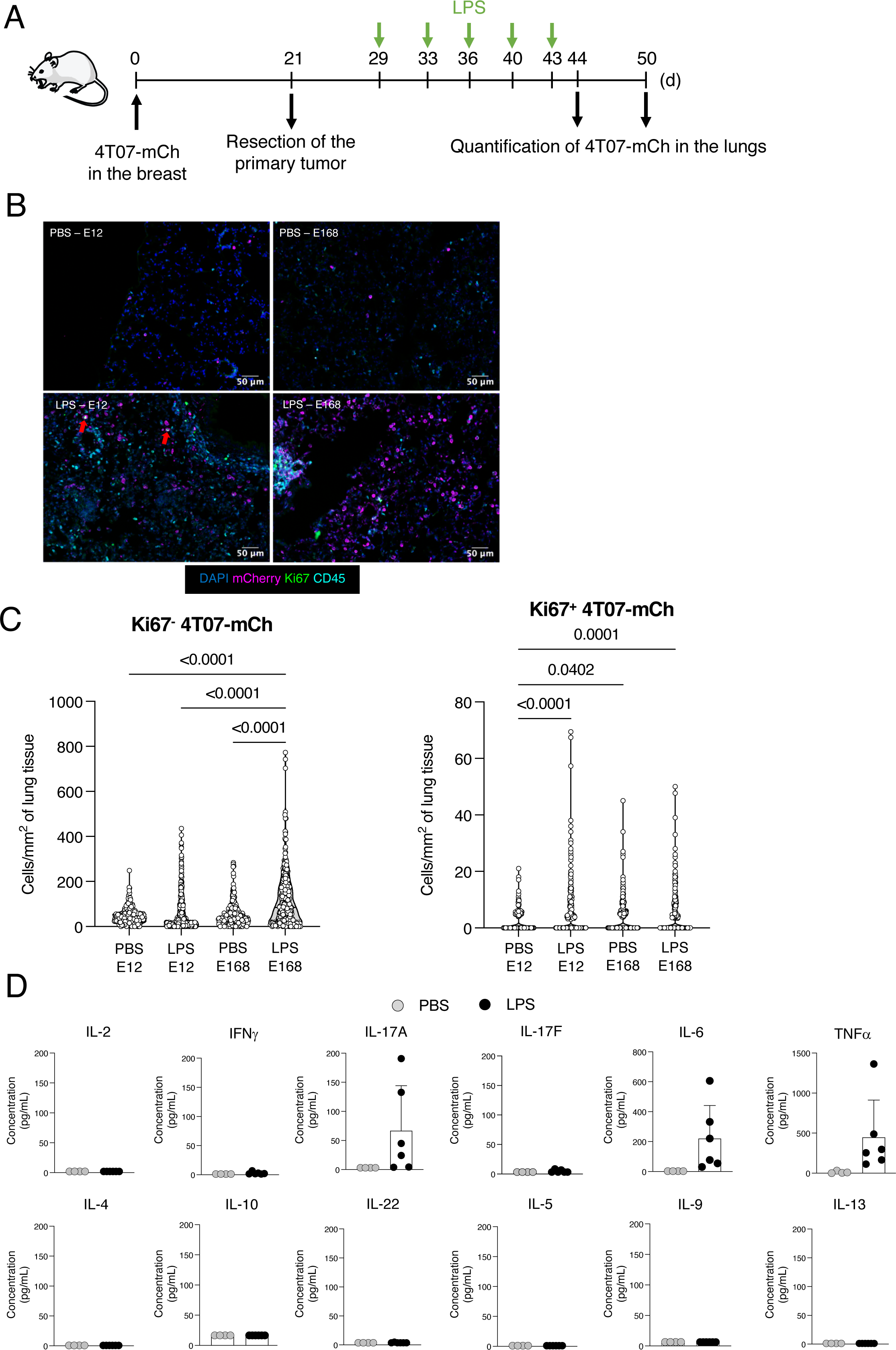
LPS-induced inflammation awakens dormant cancer cells. **(A)** Experimental design. 4T07-mCh cells (10^5^) were injected into the mammary fat pad and the primary tumor was resected on day 21, after which mice received 10 µg of LPS or an equivalent volume of PBS intranasally twice per week. The lungs were collected 12 hours (E12) and 7 days (E168) after the last intranasal administration for analysis. PBS E12, n = 5; LPS E12, n = 6; PBS E168, n = 5; LPS E168, n = 6. **(B)** Immunofluorescence images of the lungs on endpoints E12 and E168. Proliferating cancer cells are identified with a red arrow. **(C)** Quantification of the DCCs in the lungs on endpoints E12 and E168. Each dot represents the number of cancer cells in a randomly chosen location in the lungs. Statistical analyses were performed with Brown-Forsythe and Welch ANOVA tests with a Dunnett’s T3 multiple comparisons test. **(D)** Concentration of cytokines in the bronchoalveolar lavage fluid on endpoint E12.

We reasoned that characterizing the inflammatory molecules in the lungs of LPS-treated mice may identify essential mediators for awakening. Thus, we collected the bronchoalveolar lavage fluid (BALF) on E12 for the quantification of inflammatory cytokines. We measured a higher concentration of TNFα, IL-6, and IL-17A in the BALF in LPS-treated mice than in PBS-treated mice, whereas the other cytokines remained at very low concentrations, below the detection limit (Figure 4D).

### IL-17A awakens dormant cancer cells

To identify individual, awakening factors, we focused on IL-17A. IL-17A is a cytokine produced by several immune cells, such as neutrophils and T cells, and is involved in anti-microbial and anti-fungal responses.^21^ The constant exposure of the airways to bacteria, spores, and, for smokers, to LPS (as a component of tobacco smoke)^22^ makes IL-17A a physiologically relevant cytokine in this context. Therefore, we i.v. injected encapsulated mRNA encoding IL-17A into mice one week after the resection of the primary tumor. Then, 6 days after starting the mRNA injections, we analyzed the lungs for metastatic outbreaks and quantified the immune cells in the BALF (Figure 5A). The production of IL-17A and the lung-targeting capacity of the material was confirmed *in vitro* and *in vivo* (Figures S5A and S5B). There was no difference in the frequency of immune cells in the BALF of control and IL-17A mRNA-treated mice (Figure 5B), suggesting that IL-17A *per se* does not recruit immune cells to the lungs. The lungs of mice treated with liposomes containing IL-17A-encoding mRNA contained significantly more 4T07-mCh cells than the control group (Figures 5C and 5D). Furthermore, a higher proportion of DCCs expressed Ki67 in the IL-17A-mRNA-treated group, indicating the proliferation of cancer cells (Figures 5C and 5D). These findings suggest that IL-17A awakens dormant cancer cells in the lungs.

**Figure 5.**
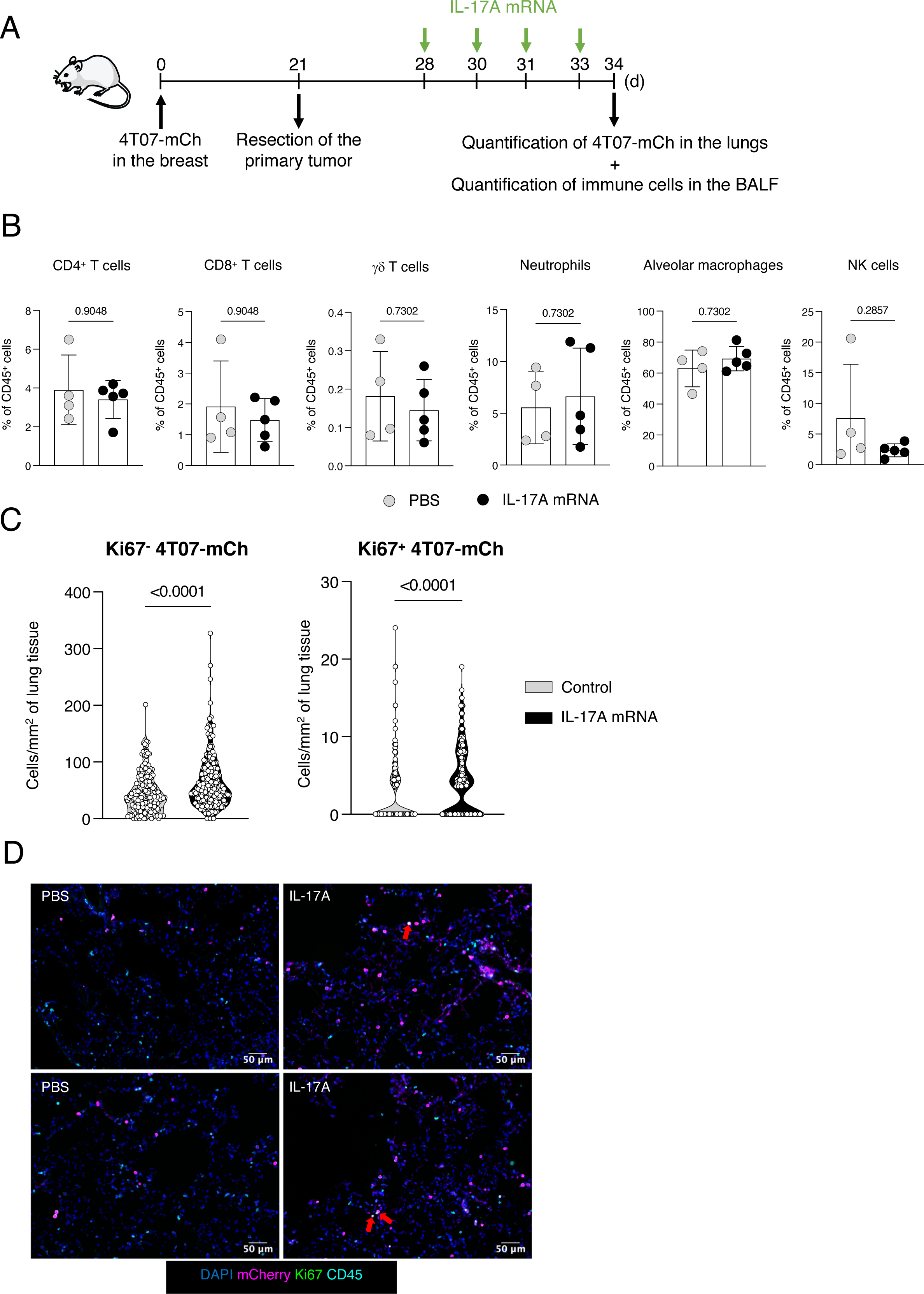
IL-17A awakens dormant cancer cells. **(A)** Experimental design. 4T07-mCh cells (10^5^) were injected into the mammary fat pad and the primary tumor was resected on day 21. One week after the resection, mice were intravenously injected with PBS or liposomes containing IL-17A mRNA. One day after the last injection, the lungs were collected for quantification of cancer cells. PBS, n = 5; DOTAP + IL-17A mRNA, n = 5. **(B)** Frequency of immune cells in the BALF on day 34. Statistical analyses were performed with a Mann-Whitney test. Each symbol represents an individual mouse. Data are shown as the mean ± standard deviation. **(C)** Quantification of Ki67^-^ DCCs and Ki67^+^ DCCs in the lungs on day 34. Each dot represents the number of cancer cells in a randomly chosen location in the lungs. Statistical analyses were performed with a Mann-Whitney test. **(D)** Immunofluorescence images of the lungs on day 34.

### LPS promotes the differentiation of T_H_17 cells in the lungs

To determine the source of IL-17A during LPS-induced inflammation in the lungs, we harvested and processed the lungs of mice that received PBS or LPS intranasally and stained the resulting single-cell suspension for IL-17A (Figure 6A). We gated on live CD45^+^ cells and, for each identified immune population, we checked for the production of IL-17A (Figure S6). At steady-state, neutrophils, CD4, CD8 and γδ T cells produced IL-17A, but LPS significantly induced IL-17A production by CD4^+^ and to a lesser extent by CD8^+^ T cells (Figure 6B). No changes in the ability of γδ T cells or neutrophils to produce IL-17A was observed after LPS administration (Figure 6B). To determine the relative contribution of different cell types to the production of IL-17A in the lungs, we gated on IL-17A^+^ live CD45^+^ cells and identified the same 4 major producers: Neutrophils, TCRαβ CD4^+^, TCRαβ CD8^+^, TCRγ8 T cells, as well as other cells (Figure S7). In steady state, neutrophils were responsible for 70% of the total IL-17A production, while TCRγ8 T and TCRαβ CD4^+^ T cells accounted for approximately 6% each (Figure 6C). However, LPS administration drastically increased the relative contribution of TCRαβ CD4^+^ T cells to the IL-17A-producing pool at the expense of neutrophils and other cell types (Figure 6C). This finding prompted us to characterize the phenotype of CD4^+^ T cells by high-dimensional flow cytometry. We identified a total of 8 clusters, with a significant increase in the frequency of the subsets expressing RORγt, consistent with a T_H_17 phenotype (Figure 6D). Of these, cluster 3 had the highest expression of IL-17A, and this subset was significantly enriched in the LPS condition (Figures 6D and 6E). These data indicate that LPS promotes the differentiation of CD4^+^ T cells into a T_H_17 phenotype and triggers their production of IL-17A.

**Figure 6.**
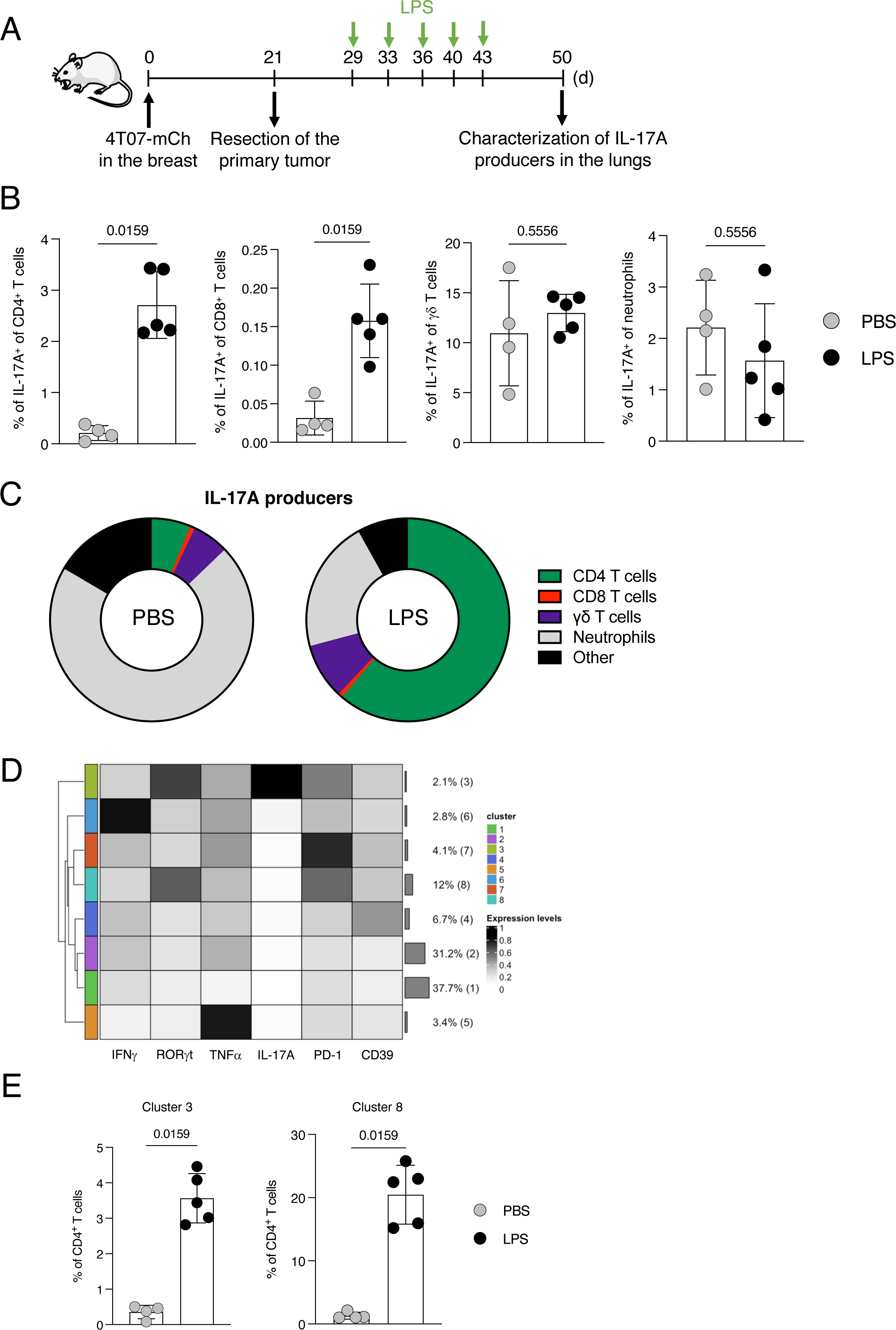
LPS induces a T_H_17 phenotype in CD4^+^ T cells. **(A)** Experimental design. 4T07-mCh cells (10^5^) were injected into the mammary fat pad and the primary tumor was resected on day 21. Mice then received 10 µg of LPS or an equivalent volume of PBS intranasally twice per week. The lungs were collected 7 days after the last intranasal administration for characterization of IL-17A-producing immune cells by flow cytometry. PBS, n = 4; LPS, n = 5. **(B)** Frequency of T cells and neutrophils that produce IL-17A. Statistical analyses were performed with a Mann-Whitney test. Each symbol represents an individual mouse. Data are shown as the mean ± standard deviation. **(C)** Proportion of IL-17A-producing cells in the lungs on day 50 after PBS or LPS admninstrations. **(D)** Heatmap with the relative expression of the tested markers. **(E)** Clusters with high expression of RORγt. Statistical analyses were performed with a Mann-Whitney test. Each symbol represents an individual mouse. Data are shown as the mean ± standard deviation.

## DISCUSSION

We aimed to identify the mechanisms involved in the different stages of metastatic dormancy, namely induction, maintenance and awakening. We previously showed that CD8^+^ T cells induce metastatic dormancy via the production of IFNγ and TNFα,^18^ but the role of CD4^+^ T cells was not addressed. Here, we show that CD4^+^ T cells are indispensable for the induction of metastatic dormancy. Our results suggest that delivery of help to CD8^+^ T cells is the major function of CD4^+^ T cells in this context, which is in line with previously published data.^23^ It is likely, however, that CD4^+^ T cells are an additional source of IFNγ. This pleiotropic cytokine acts on hematopoietic and non-hematopoietic cells,^24–28^ and is an important regulator of immune defense and homeostasis. In addition, IFNγ has direct tumor-restricting effects on cancer cells.^29–32^ By using cancer cells that can’t sense IFNγ, we found that the induction of dormancy most likely requires IFNγ sensing by cancer cells, and not by other cells.

Our observation that dormant DCCs are Ki67-negative suggests that metastatic dormancy is attributed to cell-cycle arrest on the cellular level rather than to an equilibrium of proliferation and being killed on the population level. Consequently, after its induction, dormancy likely is a cancer-cell-intrinsic feature. According to this hypothesis, depletion of NK or T cells should not lead to metastatic outbreaks. Indeed, depletion of these immune cells did not awaken dormant DCCs, as shown by absence of Ki67-expression or cluster formation of DCCs in the lungs of depleted mice. It may be unexpected that dormant DCCs are not eliminated by effector T cells, but instead persist in the face of adaptive immunity. We explain this observation by the hypothesis that the resection of the primary tumor eliminates the major source of tumor antigens, thus leading to attrition of tumor-specific effector cells. Our hypothesis is supported by the observation that tumor resection is associated with a decreased cytolytic activity in CD8^+^ T cells.^33^ The situation for NK cells seems more controversial, because a recently published study using 4T07 showed loss of dormancy in DCCs in the liver after depletion of NK cells,^34^ whereas we didn’t observe awakening of DCCs in the lungs after NK cell depletion. A possible explanation for this discrepancy, despite using similar experimental conditions, could be the anatomical and immunological differences between the lungs and the liver. Indeed, the liver contains more NK cells than the lungs.^35,36^ The relative paucity of resident NK cells in the lungs may decrease their chance of contact with the rare dormant cancer cells.

The maintenance of dormancy seems to be a cell-autonomous state. In contrast, the awakening and resuming proliferation, is an active process requiring a stimulus. Because an inflammatory response is a likely event in barrier tissues, we focused on the pro-inflammatory stimulus provided by LPS and found that the inhalation of LPS awoke dormant DCCs. Mechanistically, we identified IL-17A as a key awakening factor. Although many cell types can produce IL-17A,^37^ our results suggest that the local differentiation of CD4^+^ T cells into T_H_17 cells is essential. IL-17A is an important protective cytokine in barrier tissues, and is produced in response to fungi,^38^ which are omnipresent in inhaled air.^39,40^ Further research is warranted to elucidate how IL-17A awakens dormant DCCs and whether this pathway is unique to IL-17A.

### Limitations of the study

We unveiled the role of CD4^+^ T cells and IFNγ in the induction of maintenance of metastatic dormancy. We also showed that LPS-induced inflammation increases the local concentration of IL-17A, produced mostly by T_H_17 CD4^+^ T cells, that awakens dormant cancer cells. This study was performed using a single mouse breast cancer cell line (4T07). The lack of other models of spontaneous metastatic dormancy precluded a validation of our findings in other experimental systems. We did not address the molecular basis of IL-17A-dependent awakening of DCCs due to lack of rigorous and at the same time feasible methodologies.

## Author contributions

PP, PTdL and MvdB conceived the experiments. PP and MvdB wrote the manuscript; PP, JP, ALC, HC, MH and VC performed experiments and collected and analyzed data. MN and HK performed part of the analyses. SP provided essential reagents. MvdB secured funding; All the authors reviewed the results and approved the final manuscript.

## Acknowledgments

This work was supported by the University Research Priority Program “Translational Cancer Research” (University of Zurich; MvdB), SKINTEGRITY.ch (University of Zurich; MvdB), the Hartmann-Müller-Foundation (PP), the Swiss National Science Foundation (310030_208145 MvdB), and the Monique-Dornonville-de-la-Cour- Foundation (MvdB).

The authors thank the personnel of the Laboratory Animal Services Center (LASC, University of Zurich) and the Zurich Integrative Rodent Physiology (ZIRP, University of Zurich) for expert animal care. We thank the Flow Cytometry Facility (FCF, University of Zurich) and the Center for Microscopy and Image Analysis (ZMB, University of Zurich) for their excellent support. We thank Conrad Wyss from the Department of Dermatology for assisting in the production of the mRNA.

## Ethical approval statement

All mouse experiments were performed according to Swiss cantonal and federal regulations on animal protection and approved by the cantonal veterinary office of Zurich under the license number 26/2021 (ZH33408).

## STAR METHODS

### RESOURCE AVAILABILITY

#### Lead contact

Further information and requests for resources and reagents should be directed to and will be fulfilled by the lead contact, Maries van den Broek.

#### Materials availability

The materials generated for this study can be provided upon reasonable request.

#### Data and code availability

- All data reported in this paper will be shared by the lead contact upon reasonable request.
- This paper does not report original code.
- Any additional information required to reanalyze the data reported in this paper is available from the lead contact upon reasonable request.

### EXPERIMENTAL MODEL DETAILS

#### Animals

Eight-to-ten-week-old female BALB/cJRj mice were purchased from Janvier (Roubaix, FR). Mice were housed under specific pathogen-free conditions in individually ventilated cages at the Laboratory Animal Services Center (LASC), University of Zurich. Mice had access to food and water *ad libitum* and were kept in a 12-hour light/dark cycle. All experiments were performed according to the Swiss cantonal and federal regulations on animal protection and approved by the Cantonal Veterinary Office Zurich under the license ZH026/2021.

#### Cell lines

4T07 cells (female sex) were cultured in Dulbecco’s Modified Eagle’s Medium (DMEM, Gibco) supplemented with 5% fetal bovine serum (FBS), 2 mM L-glutamine, 100 U/mL penicillin and 100 µg/mL streptomycin. The cells were cultured at 37°C in a humid atmosphere with 5% CO_2_.

4T07 cells were lentivirally transduced to express mCherry or luciferase-ZsGreen (4T07-mCh and 4T07-LZ, respectively), as described.^18^ The *Ifngr1* gene was deleted in 4T07-mCh and 4T07-LZ cells by CRISPR/Cas9a technology. Briefly, the plasmid pSpCas9(BB)-2A-GFP (PX458) was digested with BpiI and a DNA oligonucleotide coding for a sgRNA specific for *Ifngr1* (or not specific for any sequence of the mouse genome – non-targeting sgRNA) was annealed to the digested plasmid. The resulting plasmid was transfected into 4T07-mCh and 4T07-LZ cells with the Lipofectamine^TM^ 3000 Transfection Reagent.

All cells were expanded, frozen in aliquots and stored in the gas phase of liquid nitrogen. Only cells of early passages were used for experiments. Cells were tested negative for *Mycoplasma ssp*. by PCR analysis. Cells were also tested negative for 18 additional mouse pathogens by PCR (IMPACT II Test, IDEXX Bioanalytics).

## METHOD DETAILS

### *In vivo* procedures

One hundred thousand 4T07 cells or derivatives were orthotopically injected into the 4^th^ right mammary fat pad in 50 µL of sterile PBS. Health checks were conducted twice per week. The tumor area was measured with a digital caliper as width x length.

For intravenous (i.v.) injection, 2.5 × 10^4^ 4T07-LZ and 5 × 10^4^ T cells were injected in 100 µL sterile PBS. For the depletion of CD4^+^ cells, 500 µg anti-CD4 in PBS were intraperitoneally (i.p.) injected. For the depletion of CD8^+^ cells, 500 µg of anti-CD8 in PBS were i.p. injected. For the depletion of NK cells, 50 µg of anti-asialo-GM1 in PBS were i.p. injected. As a control, 500 µg of anti-KLH or 50 µg of rabbit IgG isotype control in PBS were i.p. injected. Antibodies were administered in 100 µL sterile PBS. Macro-metastases of 4T07-LZ in the lungs were quantified using an IVIS200 imaging system (PerkinElmer). Mice were i.p. injected with 150 mg/kg of D-Luciferin and the photon flux was measured 20 minutes later. For *ex vivo* IVIS, mice were euthanized 18 minutes after the i.p. injection of D-Luciferin and the photon flux from the dissected lungs was measured immediately after.

For the resection of the primary tumor, mice were anesthetized with 3% isoflurane and given a pre-emptive, subcutaneous injection of 0.04 mg/kg of fentanyl. The area around the tumor was shaved and disinfected, and the tumor was resected. Next, the anterior part of the wound was closed with wound clips. The posterior part of the wound was stitched to ensure mobility. The wound was disinfected and the mice were subcutaneously injected with 0.1 mg/kg of buprenorphine. Mice had *ad libitum* access to water with 10 µg/mL of buprenorphine for 48 h following surgery. Based on the tumor size at resection, mice were allocated to an experimental group. The average tumor size between experimental groups was identical after randomization. This was validated by performing an unpaired t-test or one-way ANOVA with Tukey’s multiple comparisons test (for experiments with 2 or 3 groups, respectively) and obtaining non-significant *p* values.

For the awakening experiments, mice were anesthetized with 3% isoflurane and intranasally (i.n.) received 10 µg LPS in 50 µL of sterile PBS or 50 µL of PBS as described for each experiment.

To collect bronchial alveolar lavage fluid (BALF), mice were euthanized and the lungs were resected. Lungs were inflated with 1 mL of PBS containing 2 mM EDTA and the BALF was collected by aspiration. The BALF was centrifuged at 350 *g* for 5 minutes. The concentration of cytokines in the supernatant of the BALF was measured with a LEGENDplex^TM^ kit, while the pellet from the BALF was stained for flow cytometry, as described below.

For preparation of encapsulated mRNA, 90 µL of 1 µg/µL DOTAP were mixed with 10 µL of 1 µg/µL mRNA at room temperature. The structure of the mRNA is as described by Tusup *et al*.^41^ The following mRNAs were used: IL-17A and luciferase. The resulting 100 µL of the liposomal complex were i.v. injected as described for each experiment.

### Immunofluorescence

Lungs were resected, inflated with 2 mL of 4% formaldehyde, incubated in a tube with 10 mL of 4% formaldehyde for 15 minutes at room temperature, and transferred to a 15% sucrose in PBS. After overnight incubation at 4°C, the lungs were incubated with 30% sucrose in PBS for 2 days at 4°C. The lungs were then cryoembedded in Optimal Cutting Temperature (O.C.T.) Compound using a slurry with dry ice/100% ethanol. The frozen lungs were stored at -80°C. Ten-µm-thick sections were cut using a cryotome, mounted on glass slides, dried for 1 hour at 37°C, and stored at -80°C until staining.

For immunofluorescence staining, the slides were thawed and dried at 37°C for 30 minutes and fixed with 4% formaldehyde for 10 minutes. After washing with PBS, the slides were incubated for 10 minutes with PBS containing 4% BSA and 0.01% Triton- X100 to prevent unspecific binding. The slides were then washed with PBS containing 0.05% Tween 20 and incubated overnight with primary antibodies diluted in PBS containing 1% BSA at 4°C in a humidified chamber. The following day, the slides were washed with PBS containing 0.05% Tween 20 and incubated with secondary antibodies for 1 hour at room temperature in a humidified chamber. Then, the slides were washed with PBS containing 0.05% Tween 20 and stained in 100 µL of 4’,6 diamidine-2-phenylindole (DAPI) (0.5 µg) for 5 minutes. The slides were washed with PBS containing 0.05% Tween 20 and mounted with ProlongDiamond medium. The slides were scanned using the automated multispectral microscopy system Vectra 3.0 (PerkinElmer). A slide stained with the primary antibody specific for mCherry and all the secondary antibodies was used to generate the spectral profile of autofluorescence of the lungs. The multispectral images (MSIs) of each scanned slide were randomly chosen on Phenochart based solely on the expression of DAPI, to ensure the analysis was performed in an unbiased way. An average of 45 MSIs were taken per slide. The MSIs were then analyzed with inForm by performing a spectral unmixing of the individual fluorophores, adaptive tissue segmentation, cell segmentation, and phenotyping of the different cell populations. MSIs with clear immunoprecipitation regions were discarded from analysis to prevent false positives.

### Flow cytometry of tumor/lung samples

Mice were euthanized and the primary tumors or lungs were collected in complete RPMI (RPMI supplemented with 5% fetal bovine serum, 2 mM L-glutamine and 100 U/mL penicillin and 100 µg/mL streptomycin). The tissues were cut into small pieces and digested for 45 minutes at 37°C in complete RPMI supplemented with 1 mg/mL collagenase IV and 2.6 µg/mL of DNase I on a rotating wheel. After digestion, the samples were washed with PBS, mechanically disaggregated, and filtered through 70- µm filters. The filters were washed with PBS and the samples were centrifuged at 350 *g* for 5 minutes. The pellet was resuspended in 2 mL of red blood cell lysis buffer (9 g NH_4_Cl, 1.1 g KHCO_3_, 0.37 g EDTA in 100 mL of water) for 2 minutes to remove the erythrocytes. After washing with PBS, the cells were surface-stained with 50 µL of antibody mix diluted in PBS. For intracellular staining of cytokines, cells were stimulated for 4 hours at 37°C with 100 ng/mL of phorbol 12-myristate 13-acetate (PMA), 1 µg/mL of ionomycin, 1 µg/mL Brefeldin A and 1 µg/mL monensin. Non-stimulated samples were incubated under the same conditions with Brefeldin A and monensin only. After incubation, cells were surface-stained for 30 minutes at 4°C, washed with PBS and then fixed for 45 minutes at 4°C. The cells were washed with permeabilization buffer and stained overnight at 4°C with the intracellular antibody mix in permeabilization buffer. The following day, the cells were washed with permeabilization buffer, resuspended in PBS, and acquired using a CyAn ADP9 flow cytometer (Beckman Coulter), FACS LSRII Fortessa (BD Biosciences) or Cytek^®^ Aurora (5L) (Cytek).

### Flow cytometry of *in vitro*-cultured cells

To functionally validate the knockout of the *Ifngr1* gene, 4T07-LZ*^nt^*, 4T07-LZ^Δ*Ifngr*1^, 4T07-mCh*^nt^* and 4T07-mCh^Δ*Ifngr*1^ cells were incubated with 20 ng/mL of mouse IFNγ for one day. In the following day, the cells were surface-stained for PD-L1 at 4°C for 30 minutes, and then washed with PBS. Changes in the expression of PD-L1 were measured by flow cytometry with FACS LSRII Fortessa (BD Biosciences).

To validate both the functionality of the IL-17A mRNA and of the delivery system with DOTAP, HEK-293T cells were incubated for 24 hours with 100 µL of the liposomal complex containing IL-17A mRNA. The following day, the cells were incubated for 4 hours at 37°C with 1 µg/mL Brefeldin A and 1 µg/mL monensin. After incubation, the cells were surface-stained for 30 minutes at 4°C, washed with PBS and then fixed for 45 minutes at 4°C. The cells were washed with permeabilization buffer and stained in permeabilization buffer overnight at 4°C for the detection of IL-17A. The following day, the cells were washed with permeabilization buffer, resuspended in PBS, and acquired in FACS LSRII Fortessa (BD Biosciences).

## QUANTIFICATION AND STATISTICAL ANALYSIS

Group sizes and replications are provided in the figure legends. FCS files from flow cytometry were preprocessed using FlowJo. Data from compensated cell populations of interest were exported as new FCS files and imported into R. Arcsinh transformation followed by quantile normalization (1^st^ and 99^th^ quantile as boundaries) was applied to raw marker intensities.^42,43^ Dimensionality reduction was performed using the uniform manifold approximation and projection (UMAP) algorithm.^44,45^ Unsupervised clustering was achieved using the RPhenograph algorithm.^46^ Visualizations were created using the ggplot2 R package. R version 4.1.0 and Bioconductor version 3.16 were used. All immunofluorescence slides were scanned with Vectra 3.0.5 imaging system and the resulting MSIs were analyzed with inForm 2.4.6.

All figures were plotted and statistically analyzed using GraphPad Prism version 10. Statistical tests were performed as stated in each figure legend. Significance was defined as *p* values inferior to 0.05.

---

**Supplemental Figure S1:**
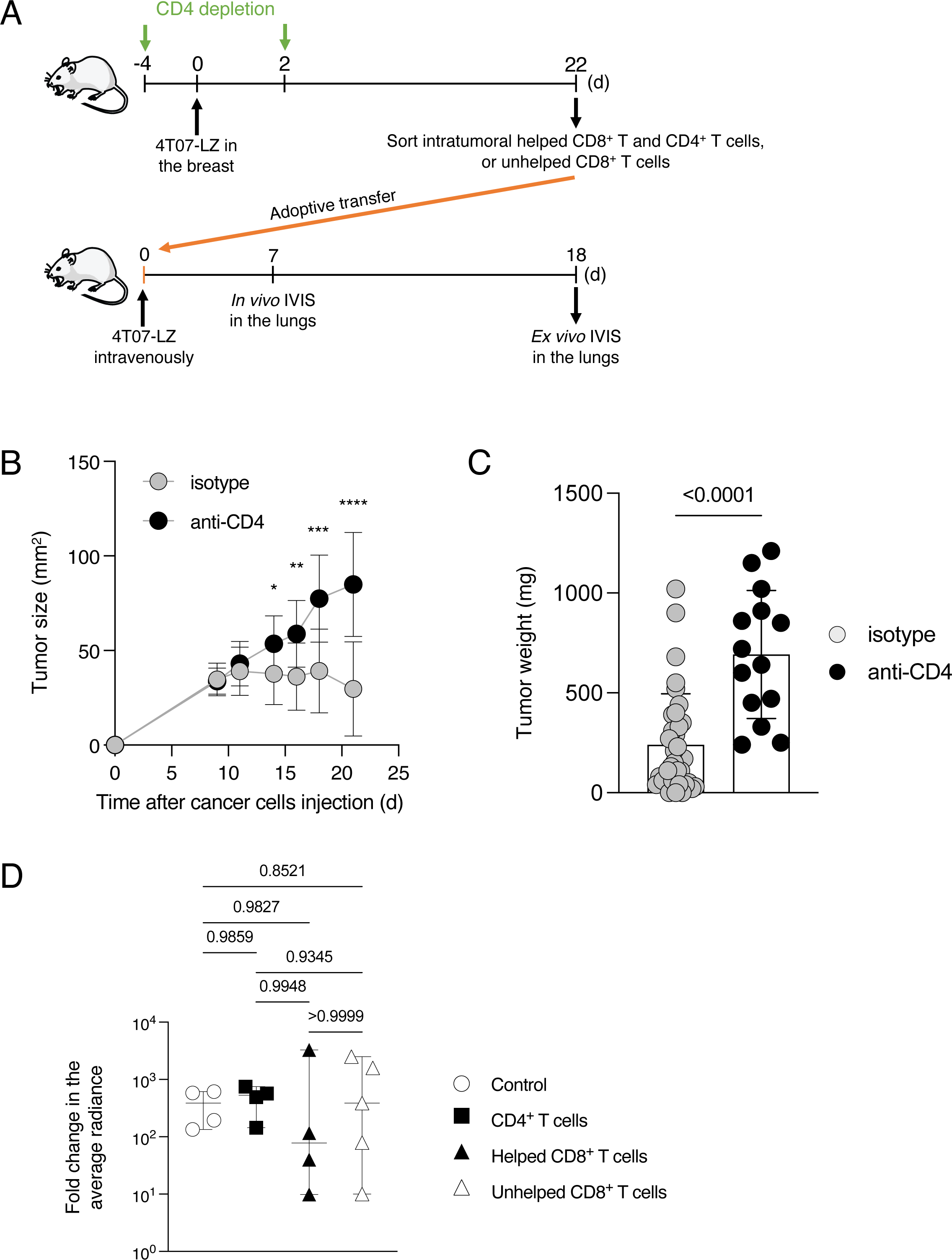
Helped CD8^+^ T cells protect against lung metastases. Related to Figure 1. **(A)** Mice received 0.5 mg isotype or anti-CD4 antibody i.p. at days -4 and +2 relative to the injection of 10^5^ 4T07-LZ cells in the mammary fat pad. The primary tumor was collected on day 22 and intratumoral CD4^+^, helped CD8^+^ or unhelped CD8^+^ T cells were isolated by fluorescence-activated cell sorting. Fifty thousand sorted T cells were i.v. injected into naïve mice, along with 2.5 × 10^4^ 4T07-LZ. The metastatic burden in the lungs was monitored by IVIS until the endpoint. **(B)** Growth curve of the primary tumor from day 9 until the endpoint. Isotype, n = 15; anti-CD4, n = 14. Statistical analyses were performed with a Mann-Whitney test. Each symbol represents the mean of the tumor size per condition. Data are shown as the mean ± standard deviation. **(C)** Tumor weight at the endpoint. The statistical analysis was performed with a Mann-Whitney test. Each symbol represents an individual mouse. Data are shown as the mean ± standard deviation. **(D)** Fold change in the average radiance of the lungs between days 7 and 18. Control, n = 4; CD4^+^ T cells, n = 4; Helped CD8^+^ T cells, n = 4; Unhelped CD8^+^ T cells, n = 5. Statistical analyses were performed with Brown-Forsythe and Welch ANOVA tests with a Dunnett’s T3 multiple comparisons test. Data is shown as median with 95% confidence interval.

**Supplemental Figure S2:**
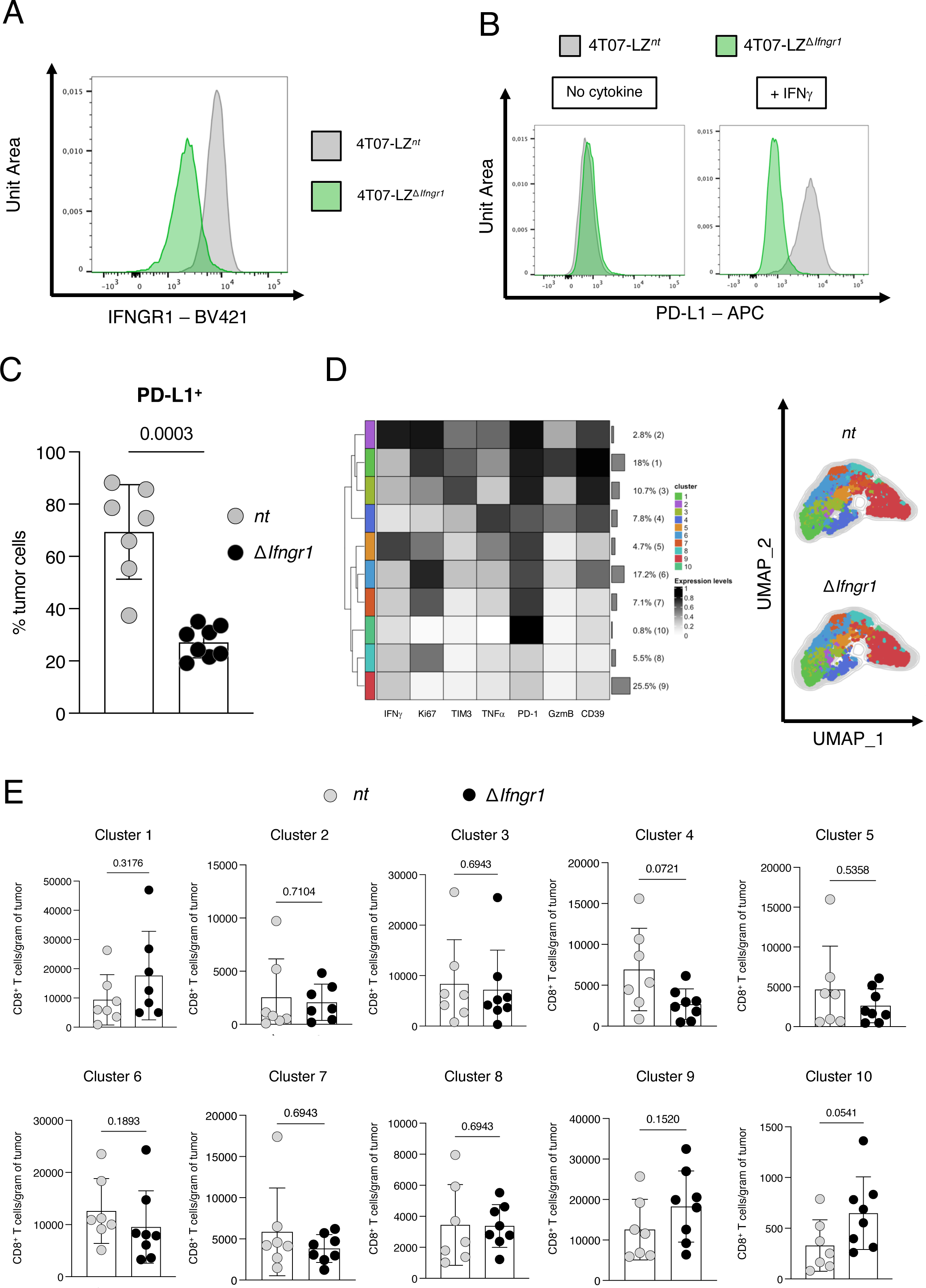
The phenotype of intratumoral CD8^+^ T cells is independent of IFNγR1 expression by cancer cells. Related to Figure 2. **(A)** Staining of IFN**γ**R1 to validate the knockout of *Ifngr1* in 4T07-mCh^Δ*Ifngr*^^1^. **(B)** Staining of PD-L1 in response to IFNg treatment in vitro, 24 hours after incubation with 20 ng/mL of IFNg. **(C)** Frequency of PD-L1^+^ cancer cells in primary tumors of 4T07-mCh*^nt^* and 4T07- mCh^Δ*Ifngr*1^ on day 14 after the injection of 10^5^ cancer cells in the mammary fat pad. The statistical analysis was performed with a Mann-Whitney test. Each symbol represents an individual mouse. Data are shown as the mean ± standard deviation. **(D)** Heatmap with the relative expression of the tested markers in the identified clusters of intratumoral CD8^+^ T cells. **(E)** Frequency of intratumoral clusters of CD8^+^ T cells. The statistical analyses were performed with a Mann-Whitney test. Each symbol represents an individual mouse. Data are shown as the mean ± standard deviation.

**Supplemental Figure S3:**
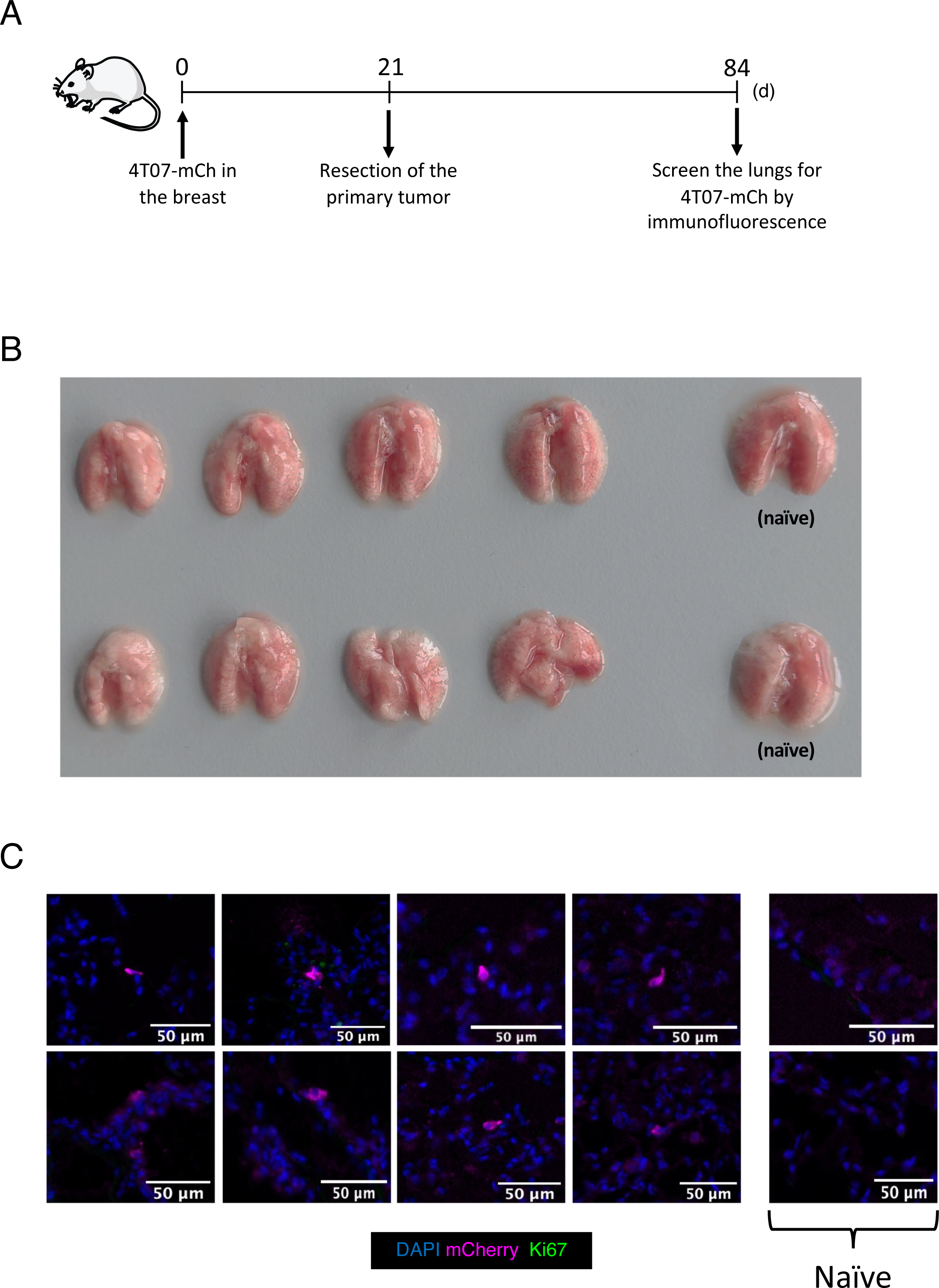
Cancer cells persist in the lungs after the resection of the primary tumor. Related to Figure 3. **(A)** Experimental design. 4T07-mCh cells (10^5^) were injected into the mammary fat pad and the primary tumor was resected on day 21. On day 84, the lungs were collected for quantification of cancer cells. Experimental group, n = 8; Naïve cage mates, n = 2. **(B)** Overview of the lungs on day 84. **(C)** Immunofluorescence images of the lungs on day 84.

**Supplemental Figure S4:**
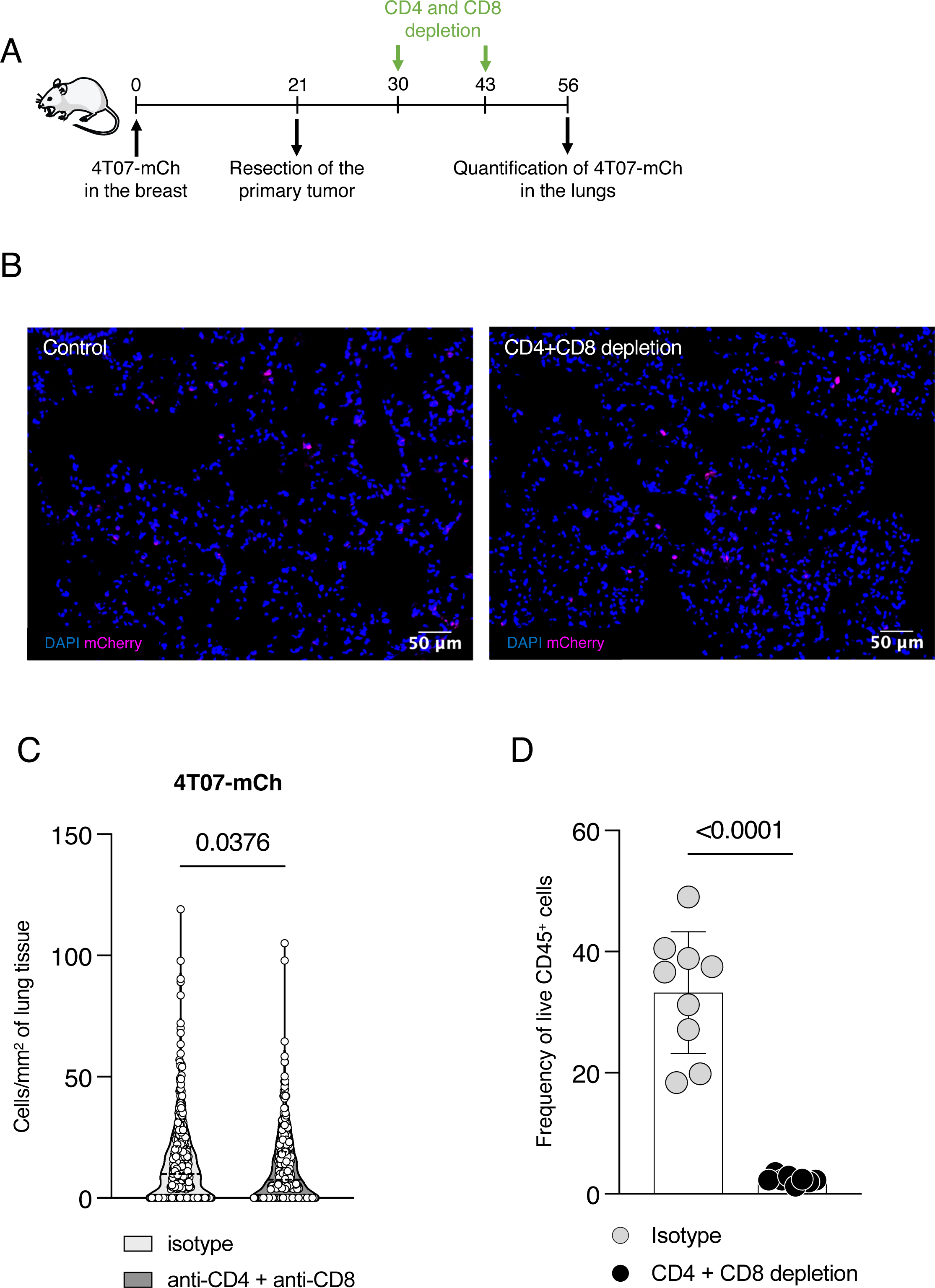
Simultaneous depletion of CD4^+^ and CD8^+^ T cells does not awaken dormant cancer cells. Related to Figure 3. **(A)** Experimental design. 4T07-mCh cells (10^5^) were injected into the mammary fat pad and the primary tumor was resected on day 21. Mice received 0.5 mg isotype or anti-CD4 antibody and anti-CD8 i.p. at days 30 and 43. On day 56, the lungs were collected for quantification of cancer cells. Isotype, n = 9; Double depletion, n = 8. **(B)** Immunofluorescence images of the lungs on day 56. **(C)** Quantification of the DCCs in the lungs on day 56. Statistical analysis was performed with a Mann-Whitney test. Each dot represents the number of cancer cells in a randomly chosen location in the lungs. The quartiles are represented by the dotted lines, the median is represented by the dashed line. **(D)** T cell depletion in blood and tumor tissue is confirmed by flow cytometry. Statistical analysis was performed with a Mann-Whitney test. Each symbol represents an individual mouse. Data are shown as the mean ± standard deviation.

**Supplemental Figure S5:**
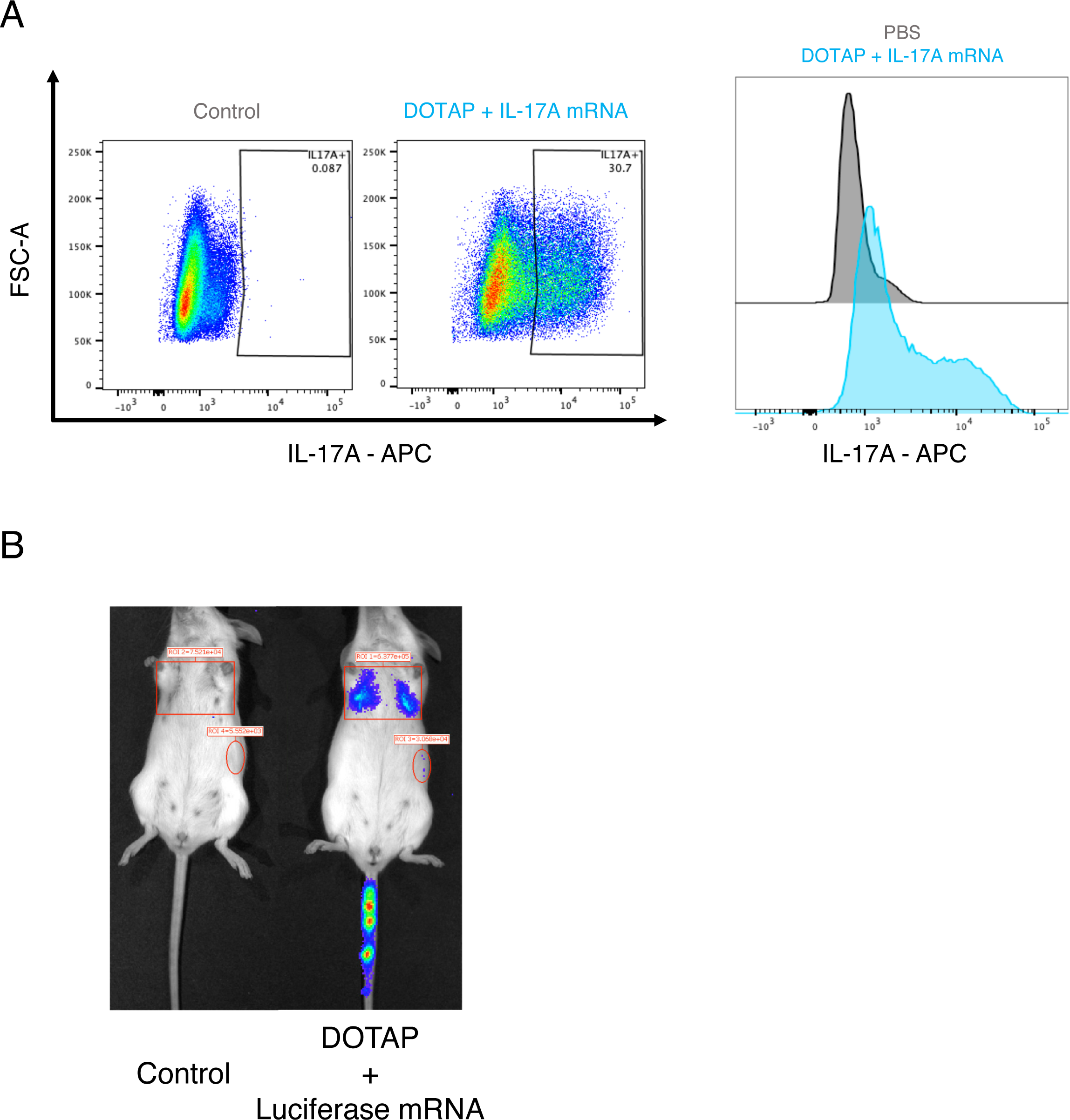
Validation of the mRNA delivery system in vitro and in vivo. Related to Figure 5. **(A)** Staining of IL-17A in vitro, 24 hours after incubation of HEK-293T with PBS or liposomes containing 10 µg of IL-17A mRNA. **(B)** Bioluminescence images of the lungs 24 hours after the injection of 100 µL of PBS or 100 µL of liposomes containing 10 µg of luciferase mRNA.

**Supplemental Figure S6:**
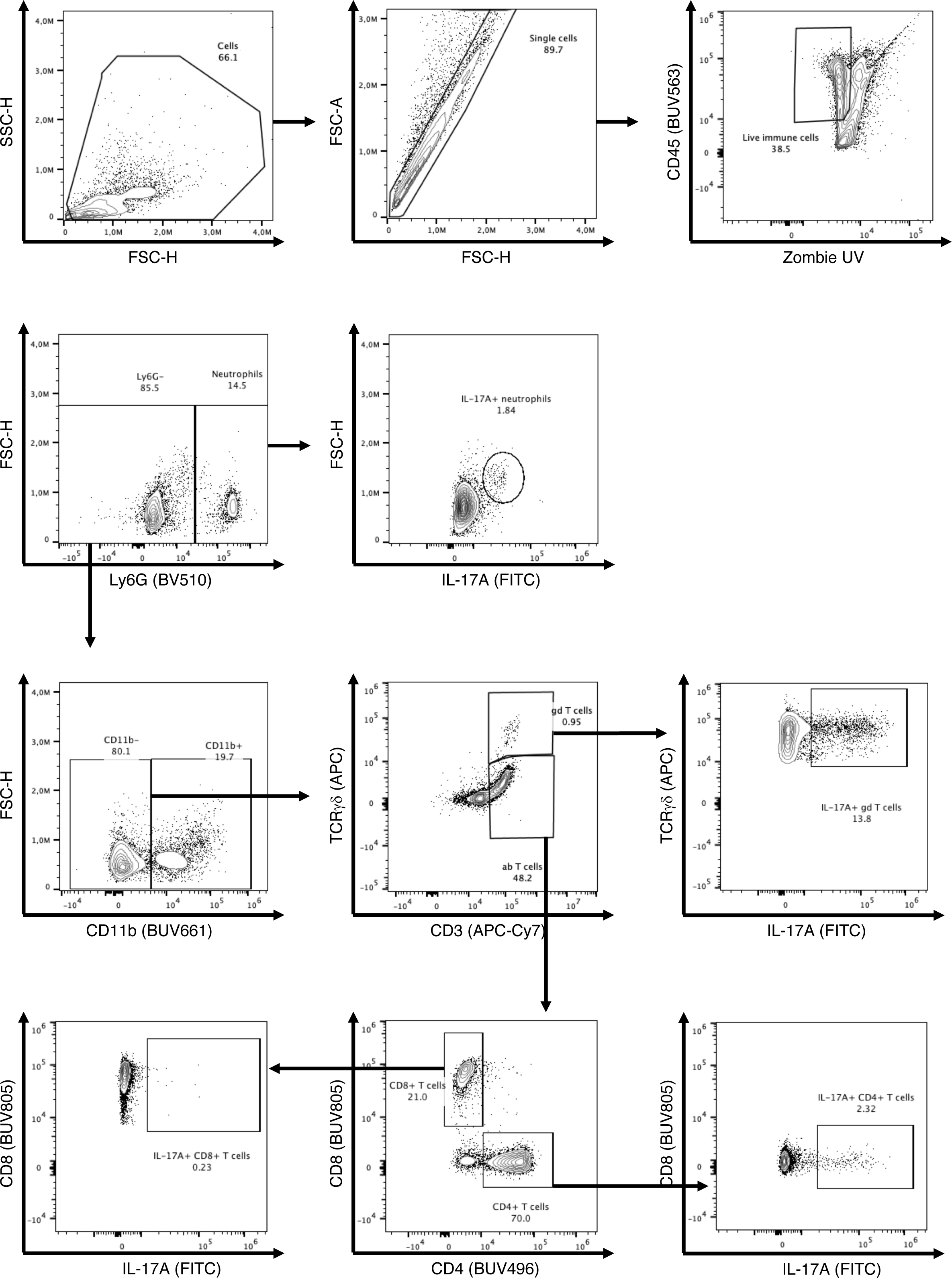
Gating strategy to identify IL-17A^+^ immune cells in the lungs. Related to Figure 6. Gating strategy to identify IL-17A-producing immune cells in the lungs on endpoint E168.

**Supplemental Figure S7:**
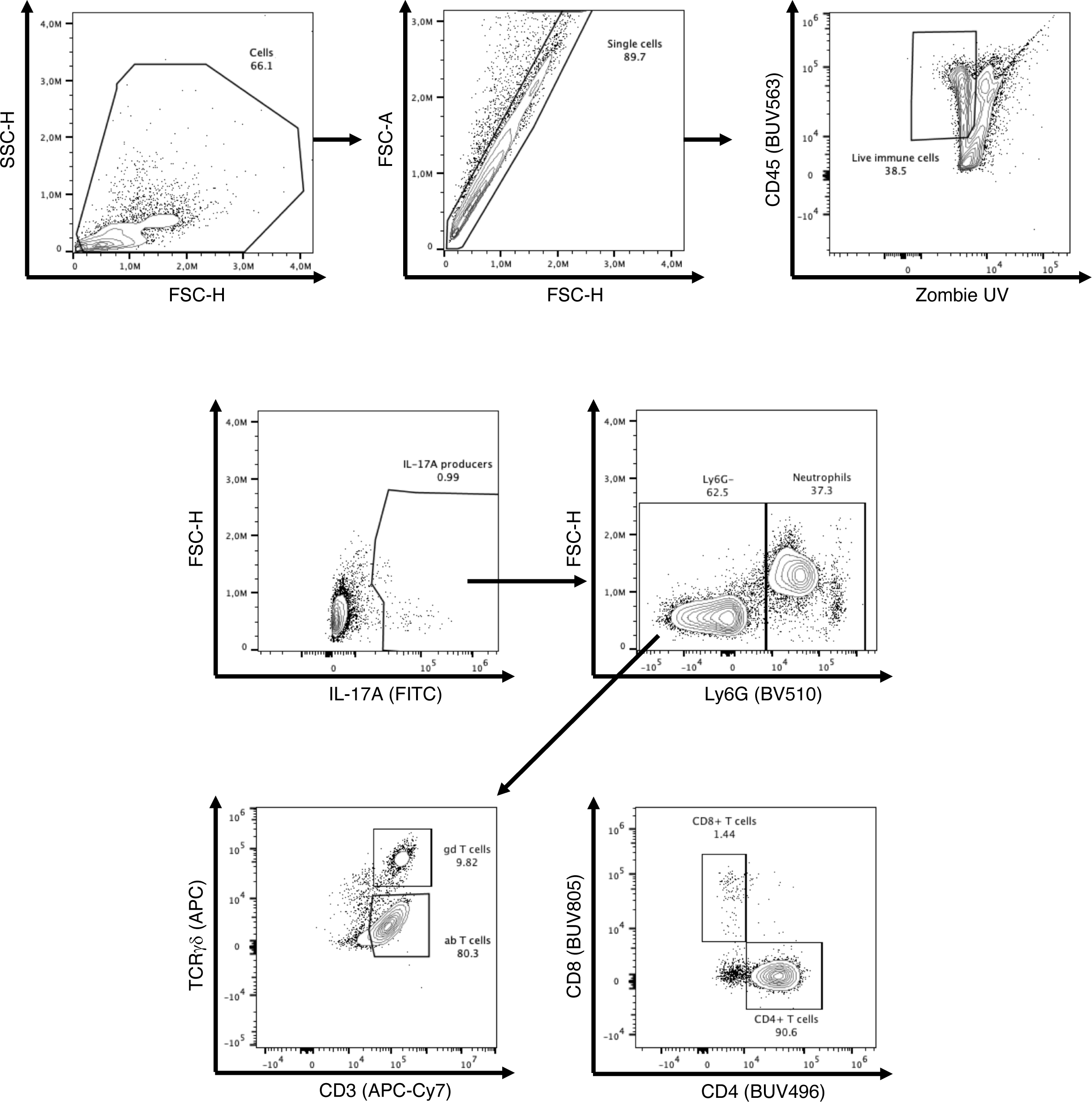
Gating strategy to determine the frequency of each immune population producing IL-17A. Related to Figure 6. Gating strategy to identify the contribution of each immune cell type in the IL-17A produced in the lungs on endpoint E168.

